# Ischemic stroke induces persistent alteration to brain stromal progenitor cells linked to chronic vascular dysfunction

**DOI:** 10.64898/2025.12.10.693193

**Authors:** Michael Candlish, Peter Breunig, Işıl Özdemir, Christina Sauerland, Stefan Günther, Kikhi Khrievono, Mario Looso, Iman Ghasemi, Katrin Schäfer, Amparo Acker-Palmer, Shamim Ashrafiyan, Johanna Ernst, Hans Worthmann, Gerrit M. Grosse, T.M Underhill, Angelos Skodras, Caroline Tscherpel, Christian Grefkes, Marcel H. Schulz, Jasmin K. Hefendehl

## Abstract

Fibrotic scar formation after stroke serves a dual role: while essential for providing structural support during post-ischemic recovery, excessive fibrosis in the chronic phase of stroke impairs regenerative processes including axonal regrowth and neovascularization. The temporal dynamics of fibrosis are critical determinants of functional outcomes, as the balance between protective scarring and regenerative capacity differs across distinct stroke phases. Consequently, strategic modulation of fibrotic processes to preserve regenerative potential represents a promising therapeutic approach in stroke recovery. To understand the cellular mechanisms underlying this fibrotic response, we investigated stromal progenitor cell composition in the post-stroke brain. The vast majority of stromal progenitor cells (SPCs) are pericytes, with minorities comprising perivascular fibroblasts (PVFs) and vascular smooth muscle cells. We demonstrate that ischemic stroke drives a long-term shift in this composition, characterized by sustained expansion of the PVF population and excessive laminin deposition in the peri-infarct region, effects that persist for at least six months post-stroke. Single-cell RNA sequencing revealed sustained transcriptional and compositional alterations in the SPC population throughout chronic post-stroke phase, driven by AP-1-mediated signaling via TNFα in both PVFs and pericytes. These changes correlate with long-term vasomotor dysfunction and capillary constriction in the peri-infarct region at six weeks post-stroke. Ischemic stroke drives aberrant, persistent PVF accumulation at the capillary bed with implications for post-stroke cerebrovascular dysfunction and recurrent stroke. Taken together, these findings reveal that ischemic stroke drives an aberrant long-term mis-localization of PVFs to the capillary bed that may have clinically-relevant implications for post-stroke cerebrovascular function as well as potential ramifications for recurrent stroke.

## Introduction

Ischemic stroke, caused by occlusion of major cerebral vessels, remains a leading cause of disability and mortality worldwide, yet therapeutic options are severely limited^1^. Treatment efficacy depends critically on rapid hospital admission for mechanical thrombectomy or intravenous recombinant tissue plasminogen activator, with effectiveness declining sharply within hours of symptom onset^2^. Consequently, clinical management predominantly focuses on limiting acute damage rather than enhancing post-stroke recovery, leaving the brain’s intrinsic repair mechanisms largely unexploited for therapeutic intervention. Central to post-stroke repair is fibrotic scar formation, which provides essential structural integrity following widespread necrosis and apoptosis within the infarct core^3,4^. However, this protective response in the long term hinders axonal regeneration and angiogenesis, creating a barrier to functional recovery^3,5^. We recently demonstrated that pericytes and perivascular fibroblasts (PVFs) constitute the brain’s resident stromal progenitor cells (SPCs), with distinct pro-angiogenic and pro-fibrotic signatures following stroke. Critically, after their activation and contribution to vascular repair, these SPCs re-localize to re-established vessels^3^. However, it remains unclear whether this re-engagement represents a return to true homeostasis or whether these cells retain long-term injury memory.

Here, we show that ischemic stroke triggers a dramatic increase in the number of PVFs within the peri-infarct region that persists until at least six months post stroke. This was associated with an accumulation of laminin close to the infarct core. Under homeostatic conditions, PVFs localize almost exclusively to primary descending arterioles^3,6,7^. Surprisingly, in the chronic post-stroke phase PDGFRα-expressing SPCs (a hallmark of PVFs) occupy both arterioles and capillaries yet exhibit pericyte-like morphology. To further understand the transcriptomic alterations taking place in SPCs in the chronic stage of stroke, we performed single cell RNA sequencing (scRNAseq) on SPCs at six weeks post stroke. Using this approach, we confirmed that these PDGFRα-expressing SPCs surrounding the infarct were in fact PVFs and not PDGFRα expressing pericytes. We further validated their identity by staining for IGFBP4, a novel PVF marker identified in our dataset, which allowed us to clearly re-identify and distinguish these PVFs from pericytes in histological sections. Analysis of the molecular mechanisms underlying these chronic SPC alterations identified the AP-1 transcriptional complex as a shared driver in both stroke-associated pericytes and PVFs, mediated specifically by TNF signaling. This persistent activation and mislocalization of PVFs was directly linked to functional vascular deficits, including chronic capillary constriction and impaired vasomotion. These data indicate that stroke triggers a lasting maladaptive shift in SPCs that is associated with changes in neurovascular integrity. Importantly, we successfully identified a circulating correlate of this specific SPC activation in the blood of stroke patients, establishing a promising new avenue for monitoring vascular health and recovery.

## Results

To investigate long-term alterations in SPCs following ischemic stroke, we performed photothrombosis on Hic1^CreERT2^;Rosa26^LSL-tdTomato^ mice (referred herein as Hic1-tdTomato), a model allowing permanent labelling of SPCs with the fluorescent reporter tdTomato upon tamoxifen administration. We then sought to differentiate perivascular fibroblasts (PVFs) from pericytes by immunohistochemical labelling for the PVF marker PDGFRα. Unexpectedly, the marked expansion of the PDGFRα^+^SPC population, initially triggered in the acute phase, was sustained until at least six months post-injury relative to controls (Fig. 1a-c). Analyzing this phenomenon in more detail, we found that PDGFRα intensity within SPCs was significantly higher in close proximity to the remaining infarct core (Fig. 1d). More intriguingly however, we found that the morphology of these cells was consistent with that of capillary pericytes rather than PVFs; the vast majority of which are found on penetrating arterioles in homeostatic conditions (Fig. 1e). Since we previously identified that PVFs in stroke play a key role in extracellular matrix protein production^3,4^, we next stained for laminin in the peri-infarct region. Consistent with the dramatic increase in PVFs in close proximity to the infarct core, we found a corresponding increase in laminin deposition around the infarct core (Fig. 1f-h). Taken together, these findings demonstrate that ischemic stroke causes long-term alterations to SPC distribution in peri-infarct tissue that is accompanied by increased deposition of ECM components, with potential ramifications for tissue repair and vascular function.

**Figure 1.**
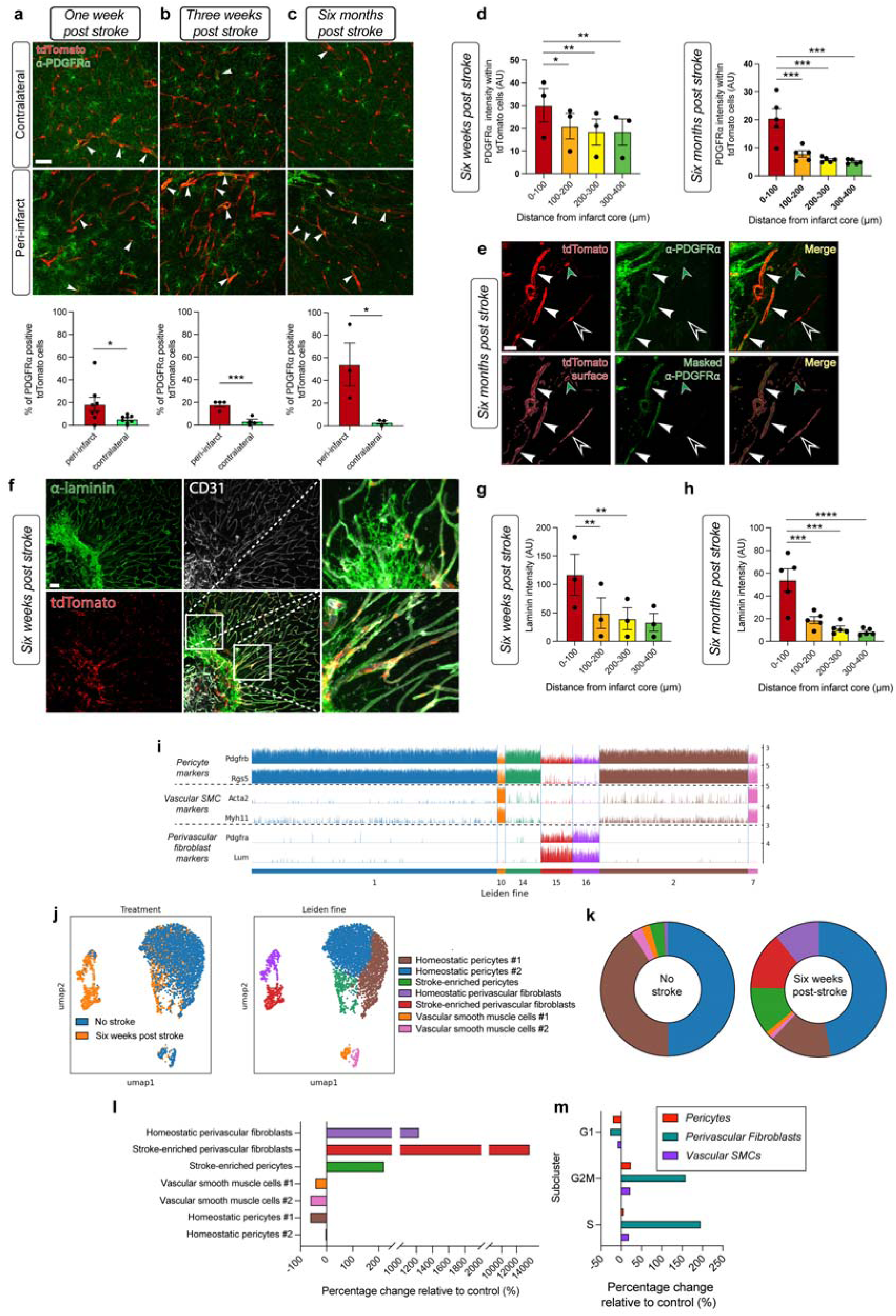
PDGFRα expression in SPCs remains elevated more than half a year after ischemic stroke. (a) The percentage of PDGFRα-expressing SPCs is significantly higher in peri-infarct tissue compared to contralateral at one week post stroke, (b) three weeks post stroke and (c) six months post stroke. Scale bar = 50 μm. (d) Histological analysis of the intensity of PDGFRα immunoreactivity within SPCs shows a significant increase proximal to the infarct core at six weeks and six months post stroke. (e) Representative image of PDGFRα expression in tdTomato^+^ cells within the peri-infarct region. PDGFRα expression is not limited to SPCs on arterioles (green arrowhead) but is found in SPCs that show pericyte morphology (white arrow heads) on capillaries (bottom left). Representative PDGFRα-negative SPC indicated with empty arrowhead. Scale bar = 30 μm. (f) Representative image of immunohistochemical labelling of laminin and CD31 (i.e. endothelial cells) at six weeks post stroke. SPCs are labelled with tdTomato. Note the abundance of extra-vascular laminin reposits. (g) Laminin intensity was significantly higher in close proximity to the infarct core at six weeks post stroke and six months post stroke. (i) Graphical representation of the expression levels of pericyte, vascular smooth muscle cell and perivascular fibroblast markers in FACS-sorted SPCs used for single cell RNA sequencing (scRNAseq) from healthy and six weeks post-stroke mice. Note that pericyte markers are highly enriched in clusters 1, 2 and 14, vascular smooth muscle cells are enriched in 7 and 10, whereas perivascular fibroblast markers were highly enriched in clusters 15 and 16. (j) UMAP of SPCs from mice six weeks post-stroke and controls (left) and UMAP illustrating the seven clusters identified using Leiden fine clustering (right). (k-l) Analysis of the percentage change relative to control of individual clusters reveals an increase not only in the “homeostatic PVF” cluster but in a subtype of perivascular fibroblast (termed “stroke-enriched PVFs) that was virtually absent in controls. Additionally, we found that while one subcluster of homeostatic pericytes (“homeostatic pericytes #1”) was substantially depleted, “homeostatic pericytes #2” remained unchanged, whereas there was a notable increase of the last remaining pericyte subcluster (termed “stroke-enriched pericytes”). (m) We identified a substantial increase in the percentage of perivascular fibroblasts in either G2M or S cell cycle stage relative to no stroke controls. (a-c) Paired two-tailed t-test. (d, e, g, h) Repeated measures one-way ANOVA with Tukey’s post hoc test. * = P < 0.05, ** = P < 0.01, *** = P < 0.001.

Given our observation of SPCs with fibroblast-like marker expression aberrantly located within the capillary bed, we sought to determine whether these cells were bona fide perivascular fibroblasts (PVFs) or pericytes that had upregulated PDGFRα following stroke. To answer this, we performed single-cell RNA sequencing (scRNA-seq) on SPCs isolated at six weeks post-stroke. We identified three major clusters corresponding to pericytes, PVFs, and a small population of vascular smooth muscle cells (vSMCs). Notably, PDGFRα expression was restricted to the PVF cluster, effectively ruling out the possibility that the PDGFRα-positive SPCs in the peri-infarct tissue were transdifferentiated pericytes^3,7^ (Fig. 1i).

Within the pericyte cluster, we identified three distinct subclusters: subcluster 2, subcluster 1, and subcluster 14. While the proportion of subcluster 1 remained relatively stable (50% of SPCs in healthy controls vs. 47% in stroke), subcluster 2 was substantially depleted in stroke (41% in controls vs. 15% in stroke), whereas subcluster 14 was markedly increased (4% in controls vs. 11% in stroke). These data suggest that the pericyte population undergoes significant and persistent remodeling well into the chronic phase. We herein refer to the depleted subcluster as ‘homeostatic pericytes #1,’ the stable subcluster as ‘homeostatic pericytes #2,’ and the expanded subcluster as ‘stroke-enriched pericytes’ (Fig. 1j-l).

Next, consistent with our immunohistochemistry results, analysis of the PVF cluster revealed a dramatic expansion in the overall proportion of PVFs (1% of SPCs in healthy controls vs. 25% in stroke) (Fig. 1j-l). Within this cluster, we identified two subclusters: subcluster 15 and subcluster 16. Remarkably, subcluster 15 was virtually absent in homeostatic conditions (0.08% of SPCs) but became the dominant PVF population after stroke (14% of total SPCs), surpassing subcluster 16 (11% of total SPCs) (Fig. 1j-l). These findings confirm not only a strong, long-lasting expansion of the PVF population but also the emergence of a novel, stroke-specific PVF subtype that is virtually absent in homeostatic conditions. From herein, we refer to cluster 16 as ‘homeostatic PVFs’ and cluster 15 as ‘stroke-enriched PVFs’ (noting that ‘homeostatic’ PVFs are also significantly expanded in stroke). Finally, regarding the vSMC cluster, both subclusters were reduced in stroke (subcluster 7: 3% in controls vs. 1% in stroke; subcluster 10: 2% in controls vs. 1% in stroke). We herein refer to cluster 10 as ‘vascular smooth muscle cells #1’ and cluster 7 as ‘vascular smooth muscle cells #2’ (Fig. 1j-l).

We next analyzed the cell cycle status of each cell type in healthy controls versus stroke conditions (Fig. 1m). We found that the cell cycle distribution remained largely unchanged in pericytes (G1: 42% vs. 33%; G2M: 29% vs. 36%; S: 29% vs. 31%) and vascular smooth muscle cells (G1: 66% vs. 61%; G2M: 14% vs. 16%; S: 20% vs. 23%). Conversely, in PVFs, we observed a substantial increase in the proportion of mitotically active cells (S-phase: 9% in controls vs. 27% in stroke; G2M: 5% in controls vs. 12% in stroke), with a corresponding decrease in the G1 fraction (86% vs. 61%). Taken together, these data indicate that six weeks after ischemic stroke, PVFs remain in the cell cycle and dramatically expand in number, accompanied by a relative reduction in the proportion of pericytes and vascular smooth muscle cells.

We then sought to identify marker genes from our scRNA-seq dataset to distinguish these SPC subtypes *ex vivo*. We found *Igfbp4* (Fig. 2a-b) to be highly selective for ‘stroke-enriched PVFs’ (subcluster 15), whereas *Ptgds* was selective for ‘homeostatic PVFs’ (subcluster 16) (Fig. 2b). *Casq2* was found to be somewhat selective for ‘homeostatic pericytes #1’ (subcluster 2), though it remained detectable in ‘homeostatic pericytes #2’ (subcluster 1) and ‘stroke-enriched pericytes’ (subcluster 14) (Fig. 2b). Intriguingly, the transcription factor *Junb*, a component of the activator protein 1 (AP-1) complex, was enriched in both ‘stroke-enriched pericytes’ and ‘stroke-enriched PVFs’. While *Junb* was also present in ‘homeostatic PVFs’, its expression appeared markedly elevated in this population under stroke conditions (Fig. 2b).

**Figure 2.**
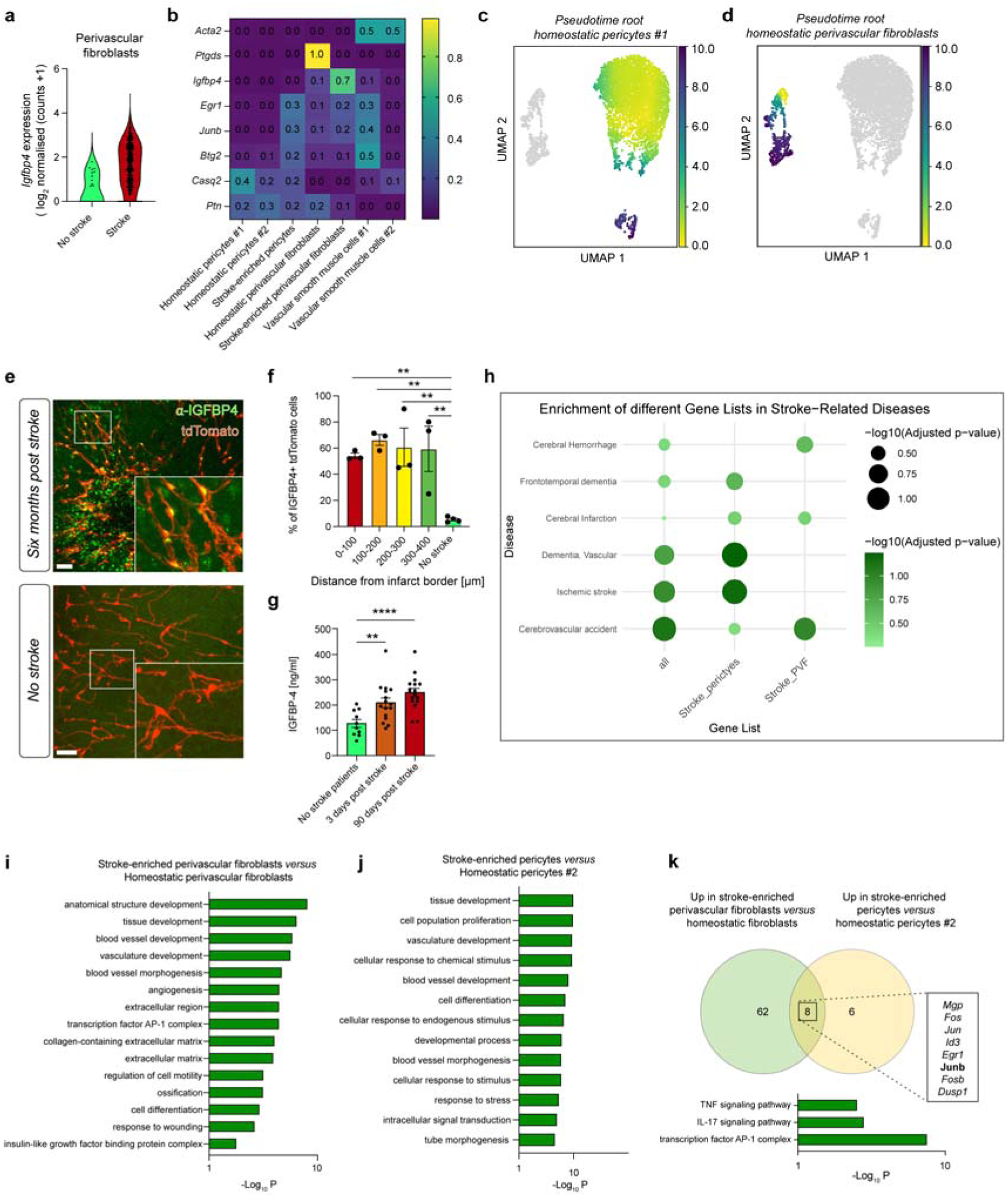
Stroke-enriched perivascular fibroblasts are readily identifiable via IGFBP4 expression. (a) *Igfbp4* is enriched in perivascular fibroblasts at six weeks post stroke. (b) Heatmap illustrating the specificity of marker genes for each SPC subcluster. (c) UMAPs showing pseudotime trajectories from homeostatic pericytes #1 and (d) homeostatic perivascular fibroblasts showing no indication of trans-differentiation of pericytes to perivascular fibroblasts. (e) Representative images of anti-IGFBP4 immunolabelling at six weeks post stroke and in no stroke mouse cortex. Scale bar = 50 μm. (f) IGFBP4 was found to be significantly higher in the peri-infarct region six weeks post-stroke compared to controls. (g) Human stroke patient blood samples contain significantly higher concentrations of IGFBP4 compared to no stroke not only in the acute phase of stroke but also the chronic phase of stroke. (h) Dot plot illustrating disease enrichment analysis of the top 200 DEGs from stroke-enriched pericytes versus homeostatic pericytes #1 and stroke-enriched PVFs versus homeostatic PVFs. (i) Bar charts illustrating the statistical significance of selected GO-terms when comparing the top upregulated genes in stroke enriched perivascular fibroblasts compared to homeostatic perivascular fibroblasts and (j) stroke-enriched perivascular pericytes compared to homeostatic pericytes #2. (k) Venn diagram showing the mutually upregulated genes in both stroke-enriched perivascular fibroblasts and stroke enriched pericytes. GO term analysis of these genes indicate that TNF and IL-17 signalling may be responsible for downstream activation of AP-1 complex in stroke SPCs. (f) Repeated measures one-way ANOVA with Tukey’s post hoc test. (g) One-way ANOVA with Dunnett’s multiple comparisons test. * = P < 0.05, ** = P < 0.01, *** = P < 0.001.

To determine whether ‘stroke-enriched PVFs’ might arise via transdifferentiation from pericytes, we performed pseudotime analysis originating from ‘homeostatic pericytes #1.’ While this trajectory suggested that vascular smooth muscle cells may originate from pericytes, none of the PVF subclusters were incorporated into the pericyte lineage trajectories, suggesting that stroke-enriched PVFs do not transdifferentiate from pericytes (Fig. 2c). We therefore performed a second pseudotime analysis originating from the ‘homeostatic PVF’ cluster. This analysis indicated that stroke-enriched PVFs are distinct from the pericyte lineage and likely originate from the pre-existing homeostatic PVF population (Fig. 2d).

To validate the cellular identity of the SPCs within the peri-infarct region, we next performed immunohistochemical labelling for IGFBP4 at six weeks post-stroke (Fig. 2e). We found that the vast majority of SPCs in this region were IGFBP4^+^, confirming their identity as the ‘stroke-enriched PVFs’ identified in our scRNA-seq dataset (Fig. 2e-f). Given that IGFBP4 is a secreted protein, we hypothesized that its levels might be elevated in the circulation of human stroke patients. Indeed, analysis of serum samples collected at three days post stroke and subsequently at 90 days post stroke revealed that IGFBP4 was significantly elevated in patients with acute ischemic stroke compared to non-stroke controls. Remarkably, IGFBP4 levels remained significantly elevated at day 90, indicating sustained PVF activation during the chronic recovery period (Fig. 2g). While IGFBP4 is markedly upregulated by activated PVFs in the post-stroke brain, its presence in circulation may also reflect contributions from other vascular cells—including vascular smooth muscle cells, endothelial cells, and systemic fibroblasts activated in response to stroke-induced systemic inflammation. Nevertheless, the strong temporal and spatial correlation between PVF activation in brain tissue and persistently elevated plasma IGFBP4 in stroke patients - from the acute phase through chronic recovery - establishes IGFBP4 as a promising biomarker of vascular remodeling and PVF-driven fibrosis after stroke.

To assess the clinical relevance of the transcriptional changes observed in stroke-enriched SPCs, we performed disease enrichment analysis using the top 200 differentially expressed genes from stroke-enriched pericytes versus homeostatic pericytes #1 and stroke-enriched PVFs versus homeostatic PVFs. We queried these gene sets against genes related to different brain diseases obtained from the DisGeNET database, which mines research articles to obtain human gene-disease associations.

Remarkably, both stroke-enriched pericyte and PVF gene signatures showed significant enrichment in multiple stroke-related disease categories (Fig. 2h). The stroke-enriched PVF gene set demonstrated particularly robust associations, with significant enrichment in cerebrovascular accident and cerebral infarction. Similarly, the stroke-enriched pericyte gene signature showed enrichment in these same disease categories, albeit with somewhat lower statistical significance. Both gene sets also showed enrichment in vascular dementia and cerebral hemorrhage, indicating broader relevance to cerebrovascular pathology. These findings validate that the persistent transcriptional alterations we observed in murine SPCs six weeks post-stroke reflect gene expression programs that are causally linked to human stroke susceptibility and cerebrovascular disease risk, underscoring the clinical relevance of our findings.

To identify molecular pathways driving the long-term alterations in SPC subpopulations, we performed gene ontology (GO) term analysis on differentially expressed genes. Comparing ‘stroke-enriched PVFs’ to ‘homeostatic PVFs,’ we identified GO terms associated with angiogenesis, extracellular matrix production, tissue development, and response to wounding (Fig. 2i). Similarly, a comparison of ‘homeostatic pericytes’ to ‘stroke-enriched pericytes’ revealed GO terms associated with blood vessel development, tissue development, and response to stress (Fig. 2j). Notably, the ‘transcription factor AP-1 complex’ was significantly enriched in both comparisons; this complex includes *Junb*, which we previously identified as a marker for both stroke-enriched pericytes and PVFs.

We next sought to identify common molecular mechanisms driving these long-term alterations by analyzing genes that were mutually upregulated in both SPC subtypes after stroke (Fig. 2k). We identified eight shared genes, and subsequent GO term analysis of this subset highlighted three key pathways: ‘TNF signaling pathway,’ ‘IL-17 signaling pathway,’ and ‘transcription factor AP-1 complex.’ These findings suggest that AP-1 complex activation via TNF or IL-17 signaling in pericytes and PVFs may be responsible for long-term SPC alterations post-stroke.

Given the evidence pointing towards AP-1 complex activation in SPCs post-stroke, we performed immunohistochemical labelling of JunB in Hic1-tdTomato mice at six weeks post-stroke. Remarkably, we found that the majority of SPCs surrounding the infarct core were positive for JunB, compared to nearly none in non-stroke controls, thus validating our scRNAseq findings (Fig. 3a-b). To determine whether TNF/IL-17 signaling is indeed responsible for the stroke-associated phenotypic changes identified via scRNAseq and histology, we performed fluorescence-activated cell sorting (FACS) to isolate SPCs. We then exposed them to TNFα or IL-17 to induce activation and subsequently treated the activated cells with the AP-1 inhibitor T5224 ^8,9^(Fig. 3c). Application of TNFα (Fig. 3d) was sufficient to drive not only an increase in cell migration (consistent with our previous findings in stroke) but additionally caused *Pdgfra* upregulation (Fig. 3e) as well as nuclear translocation of JunB (Fig. 3f). Importantly, these effects were attenuated by AP-1 inhibition thus demonstrating that TNFα is sufficient to drive AP-1 mediated phenotypic alterations in SPCs *in vitro* comparable to those identified at six weeks post stroke *in vi*vo. Contrastingly, TNFα had no significant effect on nuclear translocation of Jun, cFos or NF-κB. (Fig. 3g-i) Likewise, IL-17 treatment was insufficient to increase in cell migration, *Pdgfra* expression or JunB nuclear translocation (Supplementary Fig. 1). We next asked whether a similar phenomenon may occur in other tissues outside of the brain. To do this, we performed carotid artery injury by exposing it to FeCl_3_ to facilitate thrombosis within the vessel. Remarkably, we identified a significant increase in both (Fig. 3j-m) the percentage of PDGFRα-expressing cells and the number of JunB at 7 days post injury, suggesting that the phenomenon identified here may be relevant in vascular injury in the periphery as well as the brain.

**Figure 3.**
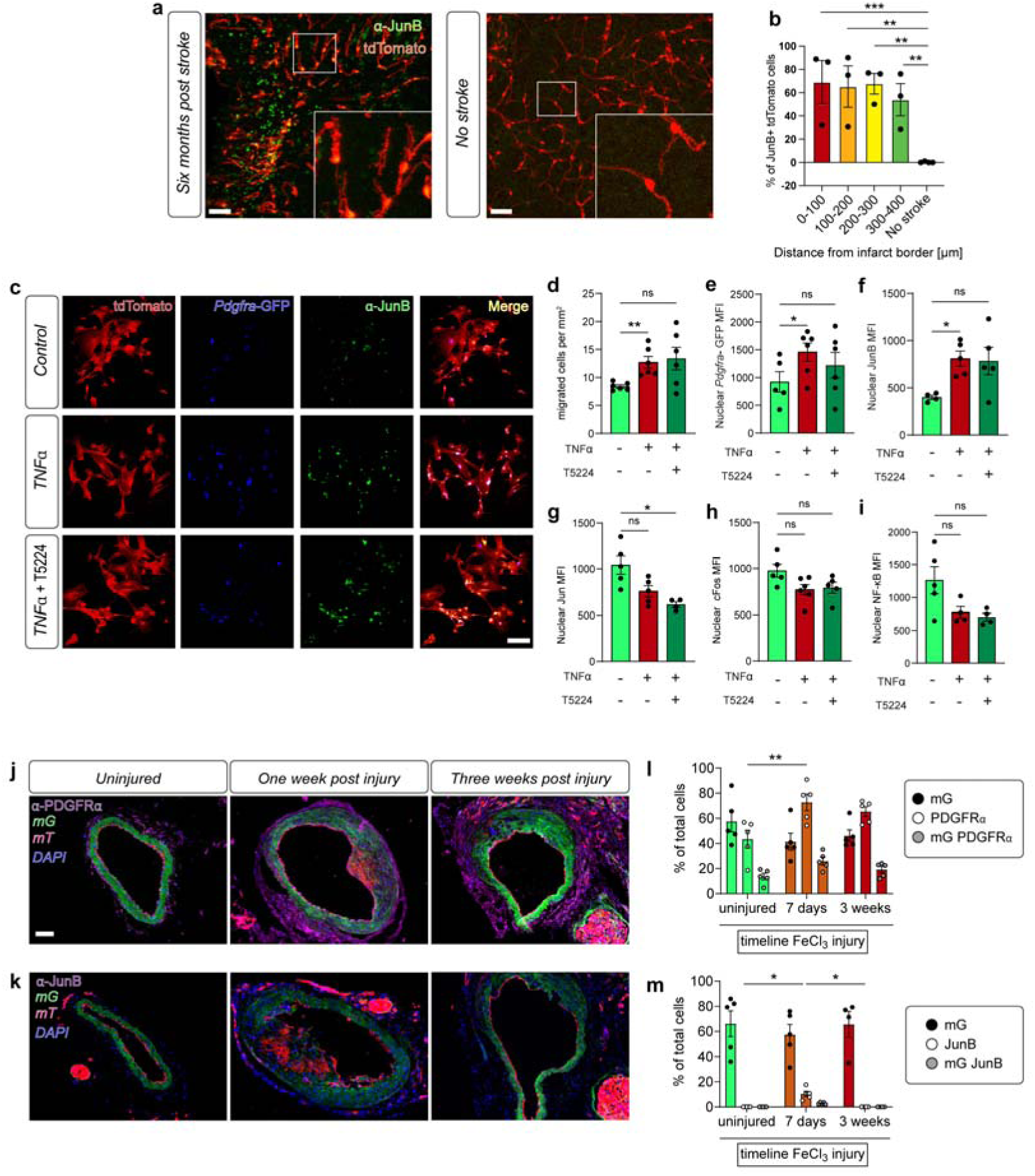
AP-1 signalling underlies phenotypic changes in SPCs post stroke. (a) Representative image of AP-1 member JunB in six weeks post-stroke peri-infarct tissue and in no stroke. Scale bar = 50 µm. (b) Percentage of JunB+ tdTomato cells is significantly higher in peri-infarct tissue compared to no stroke. (c) Representative images of immunocytochemical labelling of JunB in SPCs (expressing tdTomato and GFP via the endogenous *Pdgfra* promoter) treated with TNFα, TNFα and the AP-1 inhibitor T5224 or vehicle. Scale bar = 50 μm. Treatment with TNFα resulted in increased, (d) SPC migration, (e) increased GFP intensity (proxy for *Pdgfra* promoter activity) and (f) nuclear JunB; all of which were abolished by pre-treatment with the AP-1 inhibitor T5224. Conversely, TNFα treatment did not result in a significant increase in (g) nuclear Jun, * = P > 0.05; ** = P > 0.01 (j) Representative images of uninjured carotid artery, carotid artery one week post FeCl_3_-induced injury and carotid artery three weeks post injury in Pdgfrb-CreER^T2^ transgenic mice that were bred with membrane-Tomato (mT) membrane-Green Fluorescent Protein (mG) transgenic mice immunolabelled with anti-PDGFRα or (k) anti-JunB. Scale bar = 50 μm. (l) The percentage of PDGFRα-expressing cells was significantly increased at seven days post carotid artery injury and (m) the percentage of JunB+ cells was significantly elevated at 7 days post injury. (b) Repeated measures one-way ANOVA with Tukey’s post hoc test. (d-i). Mixed effects two-way ANOVA with Tukey’s multiple comparisons test. (l-m) ANOVA with Geisser-greenhouse correction and Dunnett’s multiple comparisons test. * = P < 0.05, ** = P < 0.01, *** = P < 0.001.

Lastly, we asked whether the pathological alterations we observed in SPCs are associated with cerebrovascular dysfunction in the chronic phase. To investigate this, we implanted cranial windows and performed *in vivo* two-photon imaging of the peri-infarct region at six weeks post-stroke (Fig. 4a). Notably, we observed vessels with stagnant flow, frequently associated with SPCs that were either mislocalized or detached from the vessel wall. Consistent with this observation, there was a significant increase in the number of occluded pial vessels compared to non-stroke controls (Fig. 4b). Additionally, quantitative analysis revealed a significantly reduced capillary diameter in the peri-infarct cortex, indicative of chronic capillary constriction (Fig. 4c).

**Figure 4.**
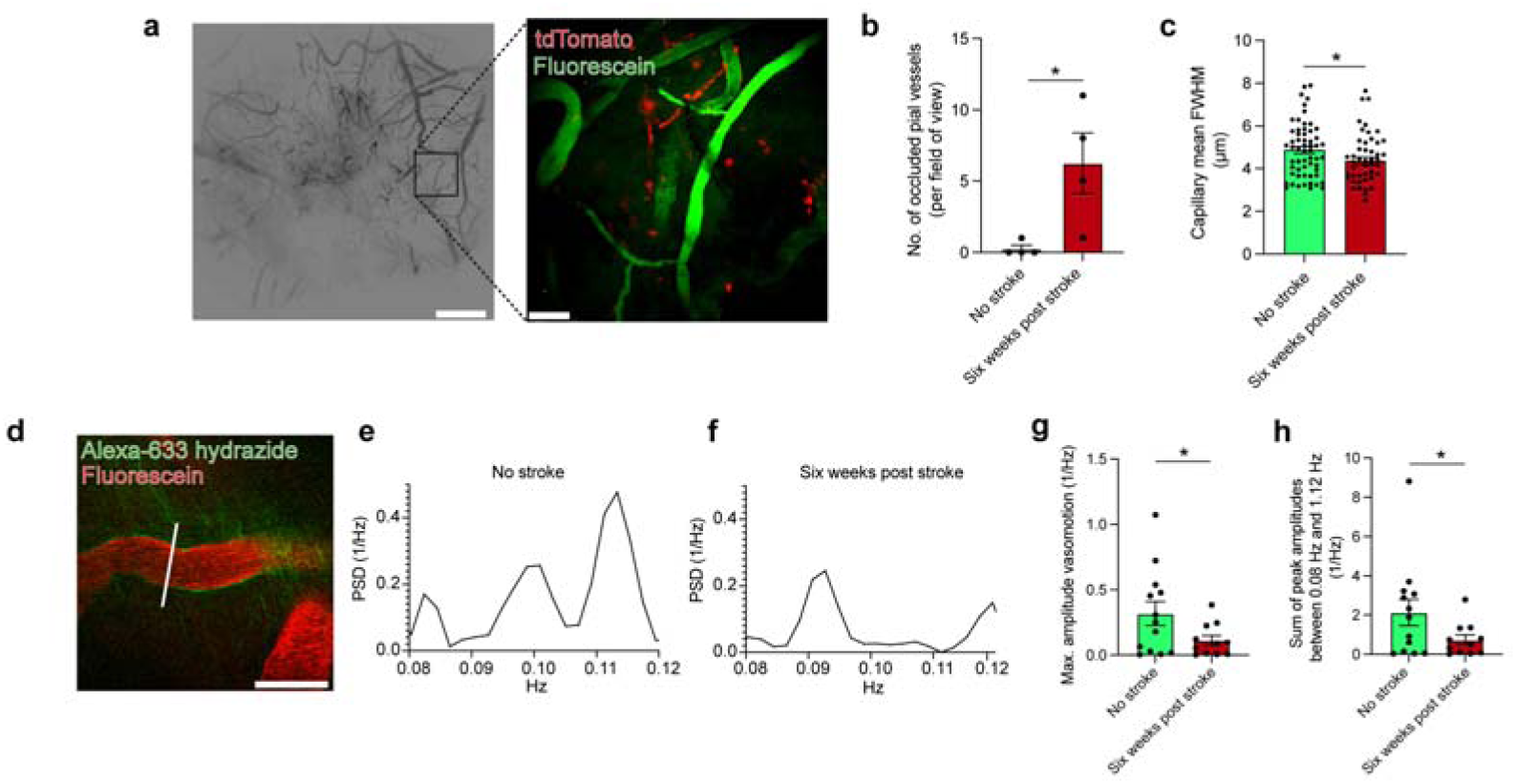
Phenotypic alterations to SPCs coincide with chronic capillary constriction and impaired vasomotion at six weeks post stroke. (a) Representative image illustrating a cranial window imaged using *in vivo* 2-photon microscopy at 5x magnification and 40x magnification in a six weeks post stroke mouse. (b) *In vivo* 2-photon imaging of mice at six weeks post stroke revealed significantly more occluded pial vessels as well as (c) chronic capillary constriction (data points represent individual capillaries) at six weeks post stroke compared to no stroke animals. (d) Representative *in vivo* 2 photon microscopy image of a raster scan targeting an alexa-633 hydrazide labelled arteriole for subsequent vasomotion analysis. (e) Representative traces showing power spectral density plots within the physiological vasomotion frequency range in no stroke mice and (f) six weeks post stroke mice. (g) Six weeks post stroke mice featured a significant reduction in the maximum amplitude as well as (h) the sum of peak amplitudes within the physiological vasomotion frequency range. Data points represent individual arterioles. (b, c, g, h) One tailed unpaired Student’s t-test. * = P < 0.05, ** = P < 0.01, *** = P < 0.001.

To further interrogate cerebrovascular function, we quantified spontaneous vasomotion in cortical arterioles (Fig. 4d-h). In the peri-infarct territory, vasomotor oscillations were markedly attenuated compared to healthy controls, manifesting as a reduced maximum amplitude of the power spectral density (PSD) as well as the sum of the peak amplitudes within the physiological vasomotion frequency range (0.08–0.12 Hz). These data demonstrate that persistent vascular dysfunction in the chronic phase extends beyond the capillary bed to the arteriolar network, compromising local perfusion and hemodynamics.

Taken together, these findings demonstrate that ischemic stroke drives long-lasting alterations in both the phenotype and demographics of SPCs, characterized by persistent AP-1-mediated transcriptional reprogramming driven by TNFα exposure. These cellular changes are associated with significant functional deficits in the peri-infarct region, including vessel occlusion, persistent capillary vasoconstriction, and impaired arteriolar vasomotion.

## Discussion

Ischemic stroke initiates a complex cascade of tissue injury and repair that extends far beyond the acute phase. While current therapeutic strategies focus predominantly on acute revascularization and neuroprotection, the long-term phase of stroke - particularly the chronic remodeling of the neurovascular unit - remain poorly understood and therapeutically unexploited^10,11^. Here, we demonstrate that ischemic stroke triggers a persistent, maladaptive reprogramming of the SPC compartment, characterized by the sustained expansion and mislocalization of PVFs to the capillary bed. This phenotypic shift is associated with a TNFα-mediated AP-1 transcriptional program and chronic vascular dysfunction, including capillary constriction and impaired vasomotion. Importantly, we show that this SPC activation signature is conserved in human stroke patients, identifying IGFBP4 as a circulating biomarker of chronic vascular remodeling.

Our observation that vascular dysfunction persists for at least six months post-stroke - a significant portion of the murine lifespan - challenges the assumption that the vasculature eventually returns to a homeostatic state after injury^10,11^. Instead, the peri-infarct region appears to enter a state of chronic "vascular frailty," characterized by stagnant flow, capillary constriction, and attenuated arteriolar vasomotion. The impairment of spontaneous vasomotion is particularly concerning, as these rhythmic oscillations are critical not only for local perfusion but also for the paravascular clearance of metabolic waste and toxic protein aggregates, such as amyloid-β^12–14^. The persistent dampening of these oscillations in the peri-infarct zone may therefore impact the brain’s ability to clear cellular debris and misfolded proteins, with possible implications for secondary neurodegeneration and vulnerability to recurrent vascular events in the same vascular bed.

Central to this pathology is the dual role of the fibrotic response. While rapid fibrosis in the acute phase is essential for sealing the blood-brain barrier and structurally isolating the necrotic core^3–5^, our data suggest that the persistence of this response is associated with maladaptive changes in the chronic phase. The accumulation of PVFs and excessive laminin deposition around capillaries likely contributes to vascular stiffening and the "no-reflow" phenomenon observed in chronic stroke^4,15^. This delineates a critical therapeutic window: interventions targeting fibrosis must preserve the beneficial acute scarring response while dampening the chronic, maladaptive expansion of PVFs. Our findings suggest that targeting PVF-specific pathways in the subacute-to-chronic phase could promote vascular recovery without compromising lesion stability.

Transcriptomically, we identified the AP-1 transcription factor complex, specifically the subunit JunB, as a regulator of this chronic activation state in both pericytes and PVFs. Our *in vitro* data provide mechanistic clarity, demonstrating that TNFα, but not IL-17, is a functional driver of this phenotype. Although GO term analysis implicated both pathways, only TNFα was sufficient to induce the complete stroke-associated signature, including migration, *Pdgfra* upregulation, and nuclear translocation of JunB. The lack of involvement of other AP-1 subunits (Jun, cFos) or NF-κB further highlights the specificity of this signaling axis. This specificity is crucial for therapeutic development, as it suggests that targeting the TNFα-JunB axis could selectively modulate the maladaptive SPC response without broadly suppressing essential inflammatory or repair mechanisms. The discrepancy between the GO analysis prediction (which flagged IL-17) and our functional validation underscores the importance of experimental verification, as pathway enrichment does not guarantee functional causality. Potential sources of chronic TNFα in the peri-infarct region include activated microglia, reactive astrocytes, and infiltrating peripheral immune cells, all of which have been shown to remain activated for months post-stroke. Furthermore, the identification of MGP (Matrix Gla Protein) and IGFBP4 as highly enriched markers in the stroke-specific PVF subset offers additional molecular handles for targeted intervention. MGP, a regulator of calcification and ECM remodeling, may play a role in the vascular stiffening observed in the chronic phase, while IGFBP4 serves as both a marker and potentially a mediator of the fibrotic response. The successful attenuation of TNFα-induced SPC activation by the AP-1 inhibitor T5224 provides proof-of-concept that this pathway is druggable and may represent a target for therapeutic intervention.

Finally, the translation of these findings to human stroke patients underscores the clinical relevance of our model. The significant and sustained elevation of plasma IGFBP4 at both day 3 and day 90 post-stroke mirrors the chronic PVF activation observed in mice. Notably, a recent study^16^ identified IGFBP4 as one of the top five proteins most strongly indicating ischemic stroke risk, suggesting that IGFBP4 may play a dual predictive role, as a marker of post-stroke vascular remodeling as well as for ischemic stroke susceptibility. While we acknowledge that circulating IGFBP4 may originate from multiple sources - including systemic fibroblasts, endothelial cells, or vascular smooth muscle cells activated by systemic inflammation - the strong temporal correlation with the chronic stroke phase suggests it reflects a genuine, sustained vascular remodeling process. Hence, IGFBP4 represents a promising candidate to monitor the extent of fibrotic remodeling and identify patients at highest risk for poor long-term recovery or recurrent stroke.

In conclusion, our study identifies a chronic, maladaptive state of the post-stroke vasculature that is associated with the aberrant persistence and mislocalization of activated PVFs. By identifying the molecular drivers and functional consequences of this response, we provide a roadmap for developing targeted therapies that extend beyond acute neuroprotection to promote long-term vascular resilience and functional recovery.

## Author contributions

M.C. and J.K.H conceived the study. M.C., P.B., I.Ö., C.S., S.G., K.K., I.G., K.S., M.L., M.H.S., S.A., and A.S., conducted experiments and/or analyzed data. J.E., H.W., G.M.G., T.M.U., C.G., C.T., contributed essential reagents. K.S., M.H.S., and A.A.P., contributed expertise and feedback. M.C., P.B., and J.K.H wrote the manuscript.

## Acknowledgements

J.K.H supported received funding from the Deutsche Forschungsgemeinschaft (DFG, German Research Foundation) IDs: 4b03813475, 419157387, Emmy Noether Award (HE 6867/3-1), SFB 1531 –Projektnummer 456687919, the Alzheimer’s Association (AARF-17-529810), and the Alzheimer Forschung Initiative e.V. (20041). BMBF (FK:01EW2308A) Neuron-ERANET, Cardio-Pulmonary Institute (CPI), EXC 2026, Project ID: 390649896, and CRC1080 (221828878). The German Centre for Cardiovascular Research (DZHK Standort Rhine Main 81Z0200101) supported A.S and M.H.S. We thank Dr. Leon Munting for helpful discussions and providing MATLAB scripts for vasomotion analysis. We thank Gisa Prange for excellent technical assistance.

## Competing interests

The authors declare no competing interests.

## Materials & Correspondence

Correspondences and requests for materials should be addressed to Prof. Jasmin Hefendehl.

## Data availability

Any additional requests for data should be addressed to Prof. Jasmin Hefendehl.

---

**Supplementary Figure 1.**
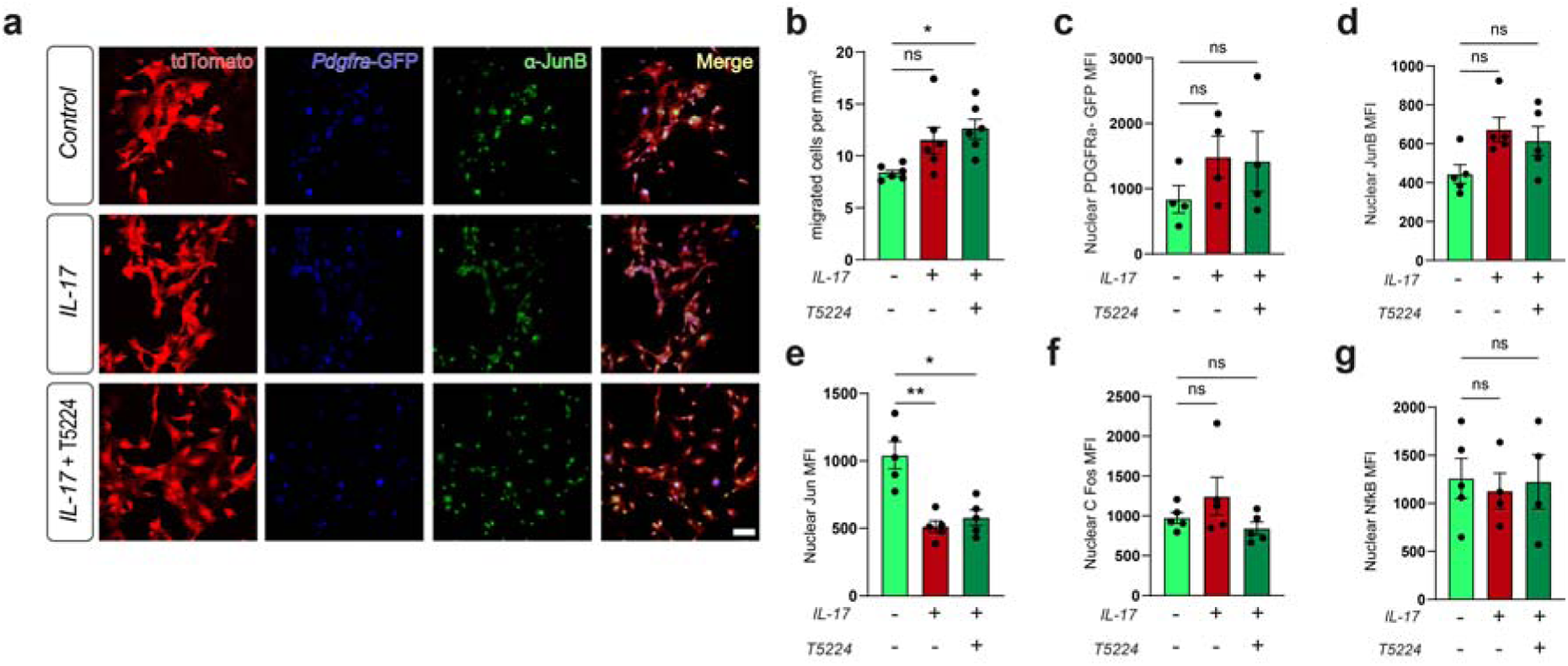
AP-1 signaling related changes in SPCs are not mediated by IL-17. (a) Representative images of immunocytochemical labelling of JunB in SPCs (expressing tdTomato and GFP via the endogenous *Pdgfra* promoter) treated with IL-17, IL-17 and the AP-1 inhibitor T5224 or vehicle. Treatment with IL-17 had no effect on (b) SPC migration, (c) GFP intensity (proxy for *Pdgfra* promoter activity) and (d) nuclear JunB, (f) nuclear cFos or (g) nuclear Nf-κB. (e) only nuclear intensity of the AP-1 member Jun was significantly reduced upon IL-17 exposure as well as after co-treatment with the AP-1 inhibitor T5224. Scale bar = 50 μm. ANOVA with Geisser-greenhouse correction and Dunnett’s multiple comparisons test. * = P < 0.05, ** = P < 0.01, *** = P < 0.001.

## Methods

### Mice (experimental animals)

Animal experiments and husbandry were carried out in accordance with guidelines established by the animal welfare committee of the Johann Wolfgang Goethe-Universität Frankfurt am Main and the Translational Animal Research Committee of the University of Mainz in accordance with regulations established by the state of Hessen (animal permit FR2022)) and the Landesuntersuchungsamt Rheinland-Pfalz (animal permit G22-1-032). Mice were kept under a standard light/dark cycle with access to food and water *ad libitum*. Hic1^CreERT2^;Rosa26^LSL-tdTomato^ knock-in mice^1^ were used for histological experiments, scRNAseq as well as *in vivo* imaging experiments. Hic1^CreERT2^;Rosa26^LSL-tdTomato^;Pdgfra-H2B-GFP knock-in mice^1^ were used for cell culture experiments. Pdgfrb-CreER^T2^ transgenic mice were cross-bred with membrane-Tomato (mT) membrane-Green Fluorescent Protein (mG) transgenic mice^2^ were used for vascular injury experiments. Induction of Cre-mediated recombination was facilitated via 5x daily i.p. injections of 100 mg/kg tamoxifen in peanut oil. All mice were kept on a C57BL/6J background.

### Ischemic stroke

Mice received an intraperitoneal (i.p,) injection of rose Bengal (Sigma) (100 mg/kg) and anesthetized using 5% isoflurane. After reaching the surgical plane of anesthesia, anesthesia was maintained at 2-3% isoflurane. The skull was exposed via a sagittal incision in the skin. The somatosensory cortex, hindlimb region was irradiated through the skull for 20 minutes with 530 nm laser light (11 mW/cm^2^) resulting in photothrombosis. Mice were administered a sub-cutaneous injection of carprofen (5 mg/kg) and the incision closed using 4/0 absorbable suture and Histoacryl. Mice were subsequently allowed to recover prior to returning to their home cages.

### Mouse model of carotid artery injury

Carotid artery injury and thrombosis was induced by applying a 10% ferric chloride (FeCl_3_) solution for 3 min onto the adventitia of the left carotid artery, as described (PMID: 11157725, PMID: 14512369, PMID: 12356640). To trace the lineage of PDGFRβ-expressing cells, Pdgfrb-CreER^T2^ transgenic mice were cross-bred with membrane-Tomato (mT) membrane-Green Fluorescent Protein (mG) transgenic mice^2^. Mice were sacrificed one and three weeks after the injury. The contralateral carotid artery was used as uninjured control. The animal experiments had been á priori approved by the Translational Animal Research Committee of the University of Mainz and the Landesuntersuchungsamt Rheinland-Pfalz (animal permit G22-1-032) and complied with national guidelines for the care and use of laboratory animals.

### Transcardial perfusion and tissue freezing for histology

Mice were euthanized with an overdose of isoflurane and, *postmortem*, transcardially perfused with 20 ml of room temperature (RT) phosphate-buffed saline (PBS) followed by 20 ml of ice-cold 4% paraformaldehyde (PFA). Brains were dissected, post-fixed for two hours in 4% PFA at 4 °C and then transferred to 30% sucrose in PBS at 4 °C for cryoprotection. Brains were then frozen in liquid nitrogen and stored at −20 °C until sectioning. 40 μm free-floating sections were prepared using a sliding microtome (Slee) and subsequently stored in freezing solution (30% glycerol, and 30% ethylene glycol in 1x PBS) at −20 °C.

### Immunohistochemistry/histology

Sections were washed 3x for 5 minutes with PBS, sequentially permeabilized and blocked with 0.5% triton X-100 in PBS and 5% donkey serum (DS) in PBS with 0.02% sodium azide for one hour each at RT. Sections were next incubated in primary antibodies (Supplementary Table 1) in PBS containing 0.5% λ-carrageenan (Sigma) and 0.02% sodium azide at 4 °C overnight. Sections were then washed 3x for 5 minutes with PBS with 0.05% tween-20 (PBST) and sequentially incubated with the appropriate secondary antibodies (Supplementary Table 1) in PBS containing 0.5% λ-carrageenan (Sigma) and 0.02% sodium azide at RT for two hours. Sections were washed 3x for 5 minutes with PBST and transferred to PBS for mounting on Superfrost Plus microscopy slides and coverslipped with Fluoromount-G.

### Wide-field and laser-scanning confocal microscopy

Brain sections were imaged at an epifluorescence microscope (Imager M.1, Zeiss) using a 10x EC Plan-NEOFLUAR 10x/0.3 objective or a laser-scanning confocal microscope (SP8, Leica) using an HC FLUOTAR L 25x/0.95 NA water-immersion objective.

### Data analysis

All statistical analyses and graph generation were performed in GraphPad Prism 9. Data analysis was performed using the Cell Counter plugin in FIJI software for the following quantifications: % of PDGFRα expressing tdTomato cells, % of IGFBP4 expressing tdTomato cells, % of JunB expressing tdTomato cells, and number of occluded vessels. Region intensity analysis of laminin and PDGFRα immunoreactivity was determined for the respective ROIs using the “measure” function of FIJI. Capillary diameter measurements were performed in FIJI using the VasoMetrics plugin^3^. The intensity of PDGFRα within SPCs was performed in Imaris software (Imaris software (v 9.7), Oxford Instruments). Figures were prepared using Fiji, Bitplane Imaris and Adobe Illustrator.

### Regions around lesion

To analyze the peri-infarct region in wide-field epifluorescence and maximum intensity projection images of SP8 confocal Z-stack images, the peri-infarct region was segmented into regions of 100 μm width starting from the border of the infarct core up to 400 μm distal from the lesion. Using FIJI software, along the border of the lesion, a ROI was manually drawn around the infarct core. This ROI was enlarged by the respective size to get regions containing all the tissue for up to 400 μm. (lesion-100 μm, lesion-200 μm, lesion-300 μm and lesion-400 μm). The ROIs were then subtracted from each other appropriately to attain the desired region ROIs (0-100 μm, 100-200 μm, 200-300 μm, and 300-400 μm). A similar approach was used for intensity analysis within SPCs in Z-stack SP8 images. A surface was generated covering the SPCs (i.e. tdTomato+), regions were then segregated into the respective areas described above based on proximity to a surface manually created over the infarct core. The mean intensity of the respective signal was then measured within SPCs in each region.

### Acute cranial window surgery and 2-photon imaging

Cranial window surgery was performed as has been previously described (REFS). Mice were anaesthetized with an i.p injection of an anesthetic cocktail of fentanyl (0.05 mg / kg), medetomidine (0.5 mg / kg) and midazolam (5 mg / kg). A 4 mm glass coverslip was then installed over the right somatosensory cortex hindlimb region. A custom-made titanium ring was subsequently attached, encircling the 4 mm glass coverslip to facilitate mounting of the mouse into the 2-photon microscope. Buprenorphine was administered for post-operative analgesia (0.8μl/g). Mice were then taken for *in vivo* 2-photon imaging. For visualization of the cerebrovasculature, an i.v. injection of 1% fluorescein-conjugated dextran (70 kDa; Sigma Aldrich 46945) in sterile saline was administered. The mouse was mounted into the microscope using a custom head-fixation device designed to perfectly articulate with the titanium ring surrounding the cranial window. Imaging was performed using a motorized custom 2-photon microscope based on the TIMAHC^4^ that is equipped with a Chameleon Ultra II laser (Coherent). Motor control and image acquisition were controlled using a MP285A (Sutter Instrument Company) and ScanImage software (MBF Bioscience). Excitation of fluorescein-conjugated dextran and alexa-633 hydrazide (ThermoFisher) was performed at 800 nm (laser power < 50 mW). Emitted light was detected using non-descanned detectors (Hamamatsu Photonics), a T560lpxr-UF2 dichroic and an ET525/50m-2p bandpass filter for fluorescein-conjugated dextran and an ET675/50m-2p for alexa-633 hydrazide. Z stack images (512×512, 1 μm step size) were taken using an A Plan-Apochromat W 40x/1.0 DIC VIS/IR (Zeiss) objective. Overview stacks were taken with a Plan-Apochromat 5x/0.16 (Zeiss) objective. Vasomotion recordings of a single optical plane were taken at 256 x 265 resolution at 2x zoom for 760 frames (frame rate 4.22 Hz).

Vasomotion analysis was performed as previous described^5^. Briefly, the acquired images were opened in FIJI and a line ROI placed perpendicularly across an Alexa-633 hydrazide positive arteriole and the ROI subsequently saved. Next, custom-made MATLAB scripts kindly provided by Dr. Leon P. Munting were used to attain the power spectral density (PSD) data from each arteriole. The output from these scripts included the maximum amplitude of the PSD within the physiological vasomotion range (0.08 – 0.12 Hz) and the sum of the amplitude peaks between 0.08 – 1.12 Hz. These data were used to discern differences in vasomotion between six weeks post stroke mice and no stroke controls.

### Statistical analysis

Statistical analyses were performed using the software Prism 9 (GraphPad Software). In general, for experiments conducted within brain regions, data was analyzed with a repeated-measure one-way ANOVA with Tukey’s multiple comparisons test unless otherwise stated. There, sphericity was assumed, and the mean of each region was compared to the mean of every other region. A mixed-effect analysis was performed if values were missing for the repeated measures analysis. Possible outliers were identified using the “Identify Outliers” function of Prism 9 using the ROUT method. Cleaned data were then further processed as described above. Unless otherwise stated, graphs display individual mice and the mean ± standard error mean.

### SPC isolation and culture

Hic1-tdTomato x PDGFRa-EGFP mice at 6-7 months of age were subjected to SPC isolation for cell culture. Mice were sacrificed and brains were collected. Following removal of the meninges, brains were minced with a scalpel and incubated with Papain (Worthington, Cat#: LK003153) for 20 minutes at 37°C in an incubator while shaking at 130rpm. Subsequently, papain was stopped using the supplied Ovomucoid protease inhibitor and tissue fragments were sequentially forced through an 18G and then 21G syringe needle for further mechanical dissociation. The resulting suspension was centrifuged for 5min at 300rcf before the myelin was removed by adding 22% BSA in sterile DPBS to the pellet and resuspending it gently. The suspension was again centrifuged, this time at 1000 rcf for 20 minutes. The resulting lipid disk was removed carefully and the Pellet washed once with DPBS. Lastly the vascular enriched pellet was resuspended in full prewarmed SPC media (ScienCell, Cat#: 1231) and seeded in 6wells. Remaining debris was washed off, and fresh media was added the day after isolation. From then on, media was changed every 2-3 days. SPCs were given a week to recover from isolation, grow to confluence and then expanded into a T25 flask. Then, tdTomato expression was triggered by *in vitro* induction via 4-Hydroxy tamoxifen (4-OHT) treatment. To this end, SPCs were exposed to 400nM 4-OHT in full SPC media for seven days. Media was replaced every day during this period. After *in vitro* induction, SPCs were detached, stained with Zombie Green^TM^ (Biolegend, Cat#: 423111, 1:200 from stock in DPBS) on ice for 15 min and FACS sorted for ZombieGreen^TM-^/tdTomato^+^ Cells using a BioRad S3e (B/Y) Cell Sorter/Analyzer. The sorted cells were brought back in tdTomato^pure^ culture, further expanded and used for experiments between passages 4-7 to ensure conservation of cellular identity *in vitro*.

### Inflammatory stimulation of cultured SPCs

SPCs were seeded into 8well object slides with detachable chambers (Sarstedt, Cat#: 83.3930.101) at 10.000 cells/well. The day after seeding, floating debris was washed off and SPCs received either a pretreatment with the selective AP-1 inhibitor T5224 (Merck, Cat#: TA9H97BAEC5E) at 20µM or DMSO for vehicle treatment. Pretreatments were conducted for 1h before either 50ng/ml TNFα, 100ng/ml IL-17 or combined TNFα (10ng/ml) and IL-17 (20ng/ml) were added to the cells. SPCs were exposed to the inflammatory stimuli for 2h and subsequently processed for Immunocytochemistry.

### Immunocytochemistry

For immunocytochemistry, media was removed and SPCs were washed once with DPBS. Next, cells were fixed with ice cold fixation buffer (4% PFA, 3% Sucrose in DPBS) for 15 minutes at room temperature. After fixations, SPCs were washed with PBS and permeabilized with 0,5% Triton-X-100 in PBS for 20 minutes at room temperature. SPCs were again washed with PBS and then blocked with 5% normal donkey-serum in PBS for 1h at room temperature. After blocking, SPCs were exposed to primary antibodies against either JunB (Cell Signaling, Cat#: 3753, diluted 1:300), Jun (Cell Signaling, Cat#: 9165, diluted 1:300), cFos (invitrogen, Cat#: PA5-143600, diluted 1:2000) or NF-κB (invitrogen, Cat#: PA5-143600, diluted 1:500) diluted in antibody buffer (0,5% λ-carrageenan and 0,02% sodium azide in PBS) over night at 4° C shaking at 70rpm. The next day, SPCs were washed three times with PBS-T before adding the appropriate (Supplementary Table 1) secondary antibodies in antibody buffer for 2h at room temperature in the dark while shaking at 70rpm. Afterwards, SPCs were washed three times with PBS-T and stained with DAPI (1µg/ml) for 10 minutes at room temperature in the dark. After DAPI staining, cells were washed two times with PBS-T and one final time with PBS before mounting. Cells were imaged using a Keyence BZ9000E epifluorescence microscope and image analysis was conducted in FIJI software.

### SPC migration assay

For migration assays, SPCs were seeded on the top side of 6well transwell inserts with 1µm pore size at 80.000 cells per membrane and left to attach overnight. The next day, cells were washed and then starved with starvation media (DMEM supplemented with 1% P/S and 0,5% FCS) for 4h. For pretreatment, starvation media contained either 20µM T5224 or DMSO for vehicle treatment. After starvation, SPCs received pre warmed full SPC media containing either 15ng/ml TNFα, 40ng/ml IL-17 or both combined (10ng/ml TNFα and 20ng/ml IL-17). Each experimental condition was prepared in duplicates. SPCs were left for migration with inflammatory stimuli for 20h. Afterwards, cells were washed with DPBS and non-migrated cells on the top side of the membranes were removed using sterile cotton swabs. Membranes were washed again twice with DPBS before fixation for 10 minutes at room temperature with fixation buffer. After fixation, membranes were washed three times with DPBS and stained with DAPI for 10 minutes at room temperature in the dark. Membranes were washed three more times with DPBS before mounting. Membranes were imaged using the 20x objective on a Keyence BZ9000E epifluorescence microscope and image analysis was conducted in FIJI software.

### Cell culture analysis

To determine the number of migrated cells, for each image a gaussian blur was applied with a radius of “r=3”. Next, a threshold was applied to each image according to Huang’s method^6^ to create a binary image. Then particles were counted in the field of view, excluding particles touching the edges. Two images were taken at random spots on each membrane with two technical replicates per condition for 6 biological replicates. From the particle count and the meta data, the number of particles/ mm^2^ were calculated by generating the mean of all images taken for each biological replicate.

For the quantification of nuclear JunB, Jun, cFos, Nf-κB and PDGFRα expression via nuclear eGFP, a nuclear mask was generated first using the DAPI channel for each image. To this end, a gaussian blur was applied with a radius of “r=3”. Next, a threshold was applied to each image according to Li’s method^7^ to create a binary image. Then, particles were measured, and regions of interest (roi) were merged to generate a nucleus mask. The nuclear signal intensity for each respective target was then measured within the nuclear mask.

### ELISA

To measure the blood levels of IGFBP-4 in either healthy controls or individuals that experiences a stroke, enzyme linked immunosorbant assay (ELISA, R&D Systems, Cat#: DY804) was used according to the supplier’s manual. Plasma of healthy individuals was kindly provided by the clinic of neurology in Frankfurt am Main, Germany. Samples of serum from stroke patients 3 days and 90 days post-stroke respectively were kindly provided by the biobank Hannover.

### FACS sorting of tdTomato^+^ SPCs

For FACS sorting, tdTomato^+^ SPCs from mouse cortex were used. Whole cortices from both hemispheres were used for healthy control animals, while for animals that experienced photothrombosis, the stroked area was punched out of the cortex and used for SPC isolation. Cerebral microvessels were isolated as described above. After myelin removal and washing, red blood cells were lysed. The pellet was resuspended in RBC lysis buffer (135mM NH_4_Cl, 10mM NaHCO_3_, 0.1mM EDTA in sterile water) and incubated on ice for 90 seconds before adding 10ml of chilled FACS AB buffer (0,5mM EDTA, 2% FCS in DPBS) followed by centrifugation. The resulting cell pellet was resuspended in FACS AB buffer. Non-viable cells were stained using DAPI and tdTomato^+^/DAPI^-^cells were sorted using a Bigfoot Spectral Cell Sorter (Thermo Fisher Scientific).

### Single cell RNA sequencing

The cell suspensions were counted with Moxi cell counter and diluted according to manufacturer’s protocol to obtain 10.000 single cell data points per sample. Each sample was run separately on a lane in Chromium controller with Chromium Next GEM Single Cell 3_ Reagent Kits v3.1 (10xGenomics). Single cell RNAseq library preparation was done using standard protocol.

Sequencing was done on Nextseq2000, and raw reads were aligned against the mouse genome (mm10) and counted by StarSolo^8^ followed by secondary analysis in Annotated Data Format. Preprocessed counts were further analyzed using Scanpy^9^. Basic cell quality control was conducted by taking the number of detected genes and mitochondrial content into consideration. We removed 6256 cells in total that did not express more than 1000 genes or had a mitochondrial content greater than 20%. Furthermore, we filtered 15476 genes if they were detected in less than 30 cells (<0.00%). Raw counts per cell were normalized to the median count over all cells and transformed into log space to stabilize variance. We initially reduced the dimensionality of the dataset using PCA, retaining 50 principal components. Subsequent steps like low-dimensional UMAP embedding^10^ and cell clustering via community detection^11^ (were based on the initial PCA. Final data visualization was done by CellxGene VIP package and GO-term analysis performed with gProfiler. In addition, stroke-related disease gene sets were obtained via literature search from the DisGenNET database for enrichment analysis. Orthologous mouse genes were mapped to human genes using the Ensembl biomart utility^12^ for comparison.

## Notes

### Competing Interest Statement

The authors have declared no competing interest.

## References

1. Saini, V., Guada, L. & Yavagal, D. R. Global Epidemiology of Stroke and Access to Acute Ischemic Stroke Interventions. Neurology 97, S6–S16 (2021).

2. Emberson, J. et al. Effect of treatment delay, age, and stroke severity on the effects of intravenous thrombolysis with alteplase for acute ischaemic stroke: a meta-analysis of individual patient data from randomised trials. The Lancet 384, 1929–1935 (2014).

3. Bernier, L.-P. et al. Brain pericytes and perivascular fibroblasts are stromal progenitors with dual functions in cerebrovascular regeneration after stroke. Nat Neurosci 28, 517–535 (2025).

4. Kelly, K. K. et al. Col1a1+ perivascular cells in the brain are a source of retinoic acid following stroke. BMC Neurosci 17, 49 (2016).

5. Protzmann, J. et al. PDGFRα inhibition reduces myofibroblast expansion in the fibrotic rim and enhances recovery after ischemic stroke. J Clin Invest 135, (2025).

6. Bonney, S. K., Sullivan, L. T., Cherry, T. J., Daneman, R. & Shih, A. Y. Distinct features of brain perivascular fibroblasts and mural cells revealed by in vivo two-photon imaging. J Cereb Blood Flow Metab 42, 966–978 (2022).

7. Vanlandewijck, M. et al. A molecular atlas of cell types and zonation in the brain vasculature. Nature 554, 475–480 (2018).

8. Ishida, M. et al. T-5224, a selective inhibitor of c-Fos/activator protein-1, improves survival by inhibiting serum high mobility group box-1 in lethal lipopolysaccharide-induced acute kidney injury model. J Intensive Care 3, 49 (2015).

9. Ye, N., Ding, Y., Wild, C., Shen, Q. & Zhou, J. Small Molecule Inhibitors Targeting Activator Protein 1 (AP-1). J. Med. Chem. 57, 6930–6948 (2014).

10. Goodman, G. W., Do, T. H., Tan, C. & Ritzel, R. M. Drivers of Chronic Pathology Following Ischemic Stroke: A Descriptive Review. Cell Mol Neurobiol 44, 7 (2023).

11. Murphy, D. P., et al. Chronic consequences of ischemic stroke: Profiling brain injury and inflammation in a mouse model with reperfusion. Physiological Reports 12, e16118 (2024).

12. Broggini, T. et al. Long-wavelength traveling waves of vasomotion modulate the perfusion of cortex. Neuron 112, 2349–2367.e8 (2024).

13. Zhang, Y.-Y. et al. High-resolution vasomotion analysis reveals novel arteriole physiological features and progressive modulation of cerebral vascular networks by stroke. J Cereb Blood Flow Metab 44, 1330–1348 (2024).

14. Murdock, M. H. et al. Multisensory gamma stimulation promotes glymphatic clearance of amyloid. Nature 627, 149–156 (2024).

15. Makihara, N. et al. Involvement of platelet-derived growth factor receptor β in fibrosis through extracellular matrix protein production after ischemic stroke. Experimental Neurology 264, 127–134 (2015).

16. Gan, X. et al. Large-Scale Plasma Proteomics Profiles for Predicting Ischemic Stroke Risk in the General Population. Stroke 56, 456–464 (2025).

## Supplementary references

1. Bernier, L.-P. et al. Brain pericytes and perivascular fibroblasts are stromal progenitors with dual functions in cerebrovascular regeneration after stroke. Nat Neurosci 28, 517–535 (2025).

2. Muzumdar, M. D., Tasic, B., Miyamichi, K., Li, L. & Luo, L. A global double-fluorescent Cre reporter mouse. Genesis 45, 593–605 (2007).

3. McDowell, K. P., Berthiaume, A.-A., Tieu, T., Hartmann, D. A. & Shih, A. Y. VasoMetrics: unbiased spatiotemporal analysis of microvascular diameter in multi-photon imaging applications. Quant Imaging Med Surg 11, 969–982 (2021).

4. Rosenegger, D. G., Tran, C. H. T., LeDue, J., Zhou, N. & Gordon, G. R. A high performance, cost-effective, open-source microscope for scanning two-photon microscopy that is modular and readily adaptable. PLoS One 9, e110475 (2014).

5. van Veluw, S. J. et al. Vasomotion as a Driving Force for Paravascular Clearance in the Awake Mouse Brain. Neuron 105, 549–561.e5 (2020).

6. Huang, L.-K. & Wang, M.-J. J. Image thresholding by minimizing the measures of fuzziness. Pattern Recognition 28, 41–51 (1995).

7. Li, C. H. & Tam, P. K. S. An iterative algorithm for minimum cross entropy thresholding. Pattern Recognition Letters 19, 771–776 (1998).

8. Dobin, A. et al. STAR: ultrafast universal RNA-seq aligner. Bioinformatics 29, 15–21 (2013).

9. Wolf, F. A., Angerer, P. & Theis, F. J. SCANPY: large-scale single-cell gene expression data analysis. Genome Biology 19, 15 (2018).

10. McInnes, L., Healy, J. & Melville, J. UMAP: Uniform Manifold Approximation and Projection for Dimension Reduction. Preprint at 10.48550/arXiv.1802.03426 (2020).

11. Traag, V., Waltman, L. & Eck, N. J. van. From Louvain to Leiden: guaranteeing well-connected communities. Sci Rep 9, 5233 (2019).

12. Dyer, S. C. et al. Ensembl 2025. Nucleic Acids Res 53, D948–D957 (2025).

